# Sex and Racial Differences in Cardiovascular Disease Risk in Patients with Atrial Fibrillation

**DOI:** 10.1101/610352

**Authors:** Wesley T. O’Neal, Aniqa B. Alam, Pratik B. Sandesara, J’Neka S. Claxton, Richard F. MacLehose, Lin Y. Chen, Lindsay G. S. Bengtson, Alanna M. Chamberlain, Faye L. Norby, Pamela L. Lutsey, Alvaro Alonso

## Abstract

**Background:** Outcomes among atrial fibrillation (AF) patients may differ according to race/ethnicity and sex due to differences in biology, the prevalence of cardiovascular risk factors, and the use and effectiveness of AF treatments. We aimed to characterize patterns of cardiovascular risk across subgroups of AF patients by sex and race/ethnicity, since doing so may provide opportunities to identify interventions. We also evaluated whether these patterns changed over time.

**Methods:** We utilized administrative claims data from the Optum Clinformatics® Datamart database from 2009 to 2015. Patients with AF with ≥6 months of enrollment prior to the first non-valvular AF diagnosis were included in the analysis. Final analysis utilized Cox proportional hazard models to estimate adjusted hazard ratios (HR) and 95% confidence intervals (CI) for cardiovascular outcomes stratified by sex and race/ethnicity. An additional analysis stratified outcomes by calendar year of AF diagnosis to evaluate changes in outcomes over time.

**Results:** In a cohort of 380,636 AF patients, women had a higher risk of ischemic stroke [HR (95% CI):1.25 (1.19, 1.31)] and lower risk of heart failure and myocardial infarction [HR (95% CI): 0.91 (0.88, 0.94) and 0.81 (0.77, 0.86), respectively)] compared to men. Black patients had elevated risk across all endpoints compared to whites, while Hispanics and Asian Americans showed no significant differences in any outcome compared to white patients. These sex and race/ethnic differences did not change over time.

**Conclusions:** We found sex and race/ethnic disparities in risk of cardiovascular outcomes among AF patients, without evidence of improvement over time.

## INTRODUCTION

Women and black patients with atrial fibrillation (AF) have been reported to have higher rates of stroke and other cardiovascular diseases compared with their male and white counterparts.^1, 2^ However, prior studies have not examined these differences across cardiovascular outcomes in sufficiently large populations. Moreover, whether these race/ethnic and sex differences have decreased or increased over time is unknown. Characterizing patterns of risk across subgroups of AF patients and their change over time can identify opportunities for intervention, leading to amelioration of existing sex and race/ethnic disparities.

The purpose of this analysis was to examine whether rates of cardiovascular outcomes among patients with AF varied by sex and race/ethnicity, and whether these rates changed over time using data from a large administrative claims database.

## METHODS

### Study population

This study used administrative claims data from the Optum Clinformatics® Datamart database between January 1, 2009 and September 30, 2015.^3^ This analysis included health plan enrollees with ≥6 months of enrollment prior to the first non-valvular AF diagnosis. AF was defined by International Classification of Diseases Ninth Revision, Clinical Modification (ICD-9-CM) codes 427.31 or 427.32 in any position on an inpatient claim or on two consecutive outpatient claims at least 7 days but less than 1 year apart, and without any inpatient diagnosis of mitral stenosis (ICD-9-CM 394.0) or mitral valve disorder (ICD-9-CM 424.0). The AF diagnosis date was defined as the earliest of the discharge date of the inpatient claim or the service date of the second outpatient claim.^4^

### Covariate assessment

Age at the time of AF diagnosis, sex (female/male), education level, and race/ethnicity were recorded at the time of health plan enrollment. Education was categorized into less than 12^th^ grade, high school diploma, less than Bachelor Degree, Bachelor Degree and beyond, and unknown. Race/ethnicity was categorized as white, black, Hispanic, and Asian American and collected from public records or imputed using commercial software (E-Tech by Ethnic Technologies). This method of imputation has been previously validated and has demonstrated 48% sensitivity and 97% specificity of classifying black race/ethnicity (relative to white).^5^ ICD-9-CM diagnosis codes in any position were used to detect the presence of comorbid conditions prior to AF diagnosis and compute CHA_2_DS_2_-VASc scores (congestive heart failure, hypertension, age ≥75 years, diabetes mellitus, stroke/transient ischemic attack, vascular disease, and age 65-75 years) for each patient.^6^ Sex was removed from CHA_2_DS_2_-VASc score calculations to account for the presence of sex as an independent variable in the model.

### Endpoint definition

Stroke, myocardial infarction, and heart failure were defined by the presence of inpatient ICD-9-CM codes in the primary position. For patients whose AF was diagnosed from inpatient claims, follow-up began the day of discharge; therefore, stroke, heart failure, or myocardial infarction occurring during the index hospitalization were not included as endpoints. All ICD-9-CM codes used for detecting study endpoints and comorbid conditions are provided in the **Supplementary Table 1**.

### Statistical analysis

Cox regression was used to estimate hazard ratios (HR) and 95% confidence intervals (CI) for each outcome by sex and race/ethnicity, adjusting for age, education, and CHA_2_DS_2_-VASc scores. Time to event was defined as days elapsed since AF diagnosis to the occurrence of the event, database disenrollment or September 30, 2015, whichever occurred earlier. The primary analysis included all eligible patients. In an additional analysis, we stratified the cohort by calendar year of AF diagnosis to evaluate changes in cardiovascular outcomes over time. For this analysis, participants were followed up for one year after AF diagnosis, with censoring occurring at the earlier of 365 days or disenrollment. Year-specific Cox regression models were used to estimate adjusted hazard ratios for each outcome by sex and race/ethnicity. Each calendar year began in January and ended in December, except for the year of 2014. Because overall follow-up ends on September 30, 2015, for the 2014 calendar year cohort we only included those diagnosed with AF January 1, 2014 through September 30, 2014. Interaction terms of calendar year of AF diagnosis with sex and race/ethnicity in the analysis including the entire sample were used to test changes in associations over time. SAS Version 9.4 (Cary, NC) was used for all analyses.

## RESULTS

A total of 380,636 participants (mean age: 73 years; 45% women; 82% white; 9% black; 7% Hispanic; 2% Asian American) were included. Table 1 provides patient characteristics at the time of AF diagnosis by race/ethnicity and sex. Compared to men, women in this cohort were more likely to have hypertension, be over the age of 75, and have had a previous stroke. Among the racial and ethnic groups, black patients were more likely to have congestive heart failure, have hypertension, and have had a previous stroke, even though they were generally younger than their counterparts. Hispanic patients were more likely to have diabetes and vascular disease.

**Table 1.** Characteristics of patients with atrial fibrillation by sex and race/ethnicity, Optum Clinformatics® 2009-2015

|  | **Men** | **Women** | **Whites** | **Blacks** | **Hispanics** | **Asian Americans** |
| --- | --- | --- | --- | --- | --- | --- |
| **N.** | 208,256 | 172,380 | 313,042 | 32,095 | 27,453 | 8,046 |
| **Age at AF Diagnosis, years\*** | 70.8 (12.3) | 75.2 (11.0) | 73.0 (11.8) | 71.0 (12.6) | 73.3 (12.4) | 73.6 (12.2) |
| **Women, %** |  |  | 44.7 | 50.0 | 46.7 | 42.7 |
| **Education, %** |  |  |  |  |  |  |
| **Less than 12<sup>th</sup> grade** | 0.6 | 0.7 | 0.2 | 0.4 | 4.9 | 1.0 |
| **High school** | 29.5 | 31.3 | 26.9 | 54.1 | 44.0 | 23.4 |
| **Less than Bachelor Degree** | 55.5 | 55.8 | 58.2 | 41.4 | 43.4 | 53.3 |
| **Bachelor Degree and beyond** | 14.1 | 12.1 | 14.4 | 3.9 | 7.4 | 22.2 |
| **Unknown** | 0.3 | 0.3 | 0.3 | 0.1 | 0.3 | 0.1 |
| **Race/ethnicity, %** |  |  |  |  |  |  |
| **Whites** | 83.1 | 81.3 |  |  |  |  |
| **Blacks** | 7.7 | 9.3 |  |  |  |  |
| **Hispanics** | 7.0 | 7.4 |  |  |  |  |
| **Asians** | 2.2 | 2.0 |  |  |  |  |
| **CHA<sub>2</sub>DS<sub>2</sub>-VASc Scores\*□** | 3.4 (2.0) | 3.9 (1.9) | 3.6 (1.9) | 3.8 (1.9) | 4.0 (2.0) | 3.7 (2.0) |
| **Congestive heart failure** | 33.1 | 34.1 | 32.3 | 41.7 | 38.5 | 31.1 |
| **Hypertension** | 80.8 | 85.6 | 82.0 | 88.9 | 87.3 | 83.1 |
| **Age ≥ 75** | 44.4 | 60.6 | 52.0 | 45.4 | 55.2 | 55.2 |
| **Age 65-74** | 28.6 | 24.1 | 26.5 | 27.9 | 25.5 | 26.5 |
| **Previous stroke** | 24.7 | 29.2 | 26.2 | 30.0 | 29.3 | 27.7 |
| **Vascular disease** | 31.5 | 31.4 | 30.7 | 34.0 | 37.8 | 30.4 |
| **Diabetes mellitus** | 36.1 | 34.4 | 33.0 | 44.8 | 48.4 | 42.6 |

\* Mean (Standard Deviation)
□ CHA<sub>2</sub>DS<sub>2</sub>-VASc without sex

Mean time to event in days for the study endpoints for all participants were as follows: stroke, 699; heart failure, 688; and myocardial infarction, 700. Heart failure was the most common outcome of the cardiovascular endpoints among all patients.

Crude rates of ischemic stroke and heart failure were higher in women than men, while men had higher rates of myocardial infarction than women (Table 2). In multivariable Cox models, however, women had a higher rate of ischemic stroke [HR (95% CI): 1.25 (1.19, 1.31)], but a lower rate of heart failure [HR (95% CI): 0.91 (0.88, 0.94)] and myocardial infarction [HR (95% CI): 0.81 (0.77, 0.86)] compared with men.

**Table 2.** Associations of sex and race/ethnicity with incidence of ischemic stroke, heart failure, and myocardial infarction in patients with atrial fibrillation, Optum Clinformatics® 2009-2015

|  | **Men** | **Women** | **Whites** | **Blacks** | **Hispanics** | **Asian Americans** |
| --- | --- | --- | --- | --- | --- | --- |
| **N.** | 208,256 | 172,380 | 313,042 | 32,095 | 27,453 | 8,046 |
| **Ischemic Stroke** |  |  |  |  |  |  |
| **Person-Years of Follow-up** | 399,921 | 328,703 | 602,989 | 55,639 | 54,431 | 15,565 |
| **N. events** | 3,251 | 3984 | 5,671 | 768 | 625 | 171 |
| **Crude IR\*** | 8.1 | 12.1 | 9.4 | 13.8 | 11.5 | 11.0 |
| **HR (95%CI)\*\*** | 1 (ref) | 1.25 (1.19, 1.31) | 1 (ref) | 1.44 (1.33, 1.55) | 1.09 (1.00, 1.19) | 1.12 (0.96, 1.30) |
| **Heart failure** |  |  |  |  |  |  |
| **Person-Years of Follow-up** | 393,421 | 323,939 | 594,354 | 54,170 | 53,442 | 15,375 |
| **N. events** | 9264 | 7994 | 13,439 | 2090 | 1422 | 307 |
| **Crude IR\*** | 23.5 | 24.7 | 22.6 | 38.6 | 26.6 | 20.0 |
| **HR (95%CI)\*\*** | 1 (ref) | 0.91 (0.88, 0.94) | 1 (ref) | 1.45 (1.38, 1.52) | 0.95 (0.90, 1.01) | 0.84 (0.75, 0.94) |
| **Myocardial infarction** |  |  |  |  |  |  |
| **Person-Years of Follow-up** | 399,723 | 330,052 | 603,902 | 55,671 | 54,586 | 15,607 |
| **N. events** | 3199 | 2386 | 4490 | 538 | 450 | 107 |
| **Crude IR\*** | 8.0 | 7.2 | 7.4 | 9.7 | 8.2 | 6.9 |
| **HR (95%CI)\*\*** | 1 (ref) | 0.81 (0.77, 0.86) | 1 (ref) | 1.12 (1.03, 1.23) | 0.95 (0.86, 1.05) | 0.89 (0.73, 1.08) |

IR, incidence rate; HR, hazard ratio; CI, confidence interval.
\*Per 1,000 person-years
\*\*Cox model adjusted for age, sex, race/ethnicity, education and CHA<sub>2</sub>DS<sub>2</sub>-VASc.

In the context of race/ethnicity, black patients presented with the highest HRs, particularly with ischemic stroke [HR (95% CI): 1.44 (1.33, 1.55)] and heart failure [HR (95% CI): 1.45 (1.38, 1.52)] (Table 2). The effect was weaker, albeit still significant, for black AF patients and their risk for myocardial infarction [HR (95% CI): 1.12 (1.03, 1.23)]. Compared to whites, Hispanic patients had a marginally higher rate of ischemic stroke, but did not differ significantly in rates of heart failure and myocardial infarction. Differences between whites and Asian Americans were not significant, save for a protective effect among Asian Americans for heart failure.

In the analysis stratified by calendar year, women had consistently higher rates of ischemic stroke (Figure 1), along with lower rates of heart failure (Figure 2) and myocardial infarction compared to men (Figure 3). Compared to white patients, black patients had consistently higher rates of stroke (Figure 4) and heart failure (Figure 5), while differences in rates of myocardial infarction over the years were small and consistent with the overall weak association between race and myocardial infarction (Figure 6). Detailed numbers for these analyses are presented in **Supplementary Tables 2-4**. There was less evidence of differences in the rates of outcomes for Hispanics compared to whites. Due to low counts within the database and for privacy protection, year-specific results for Asian Americans are not presented. Tests of interaction to determine whether sex or race/ethnicity are changing over time were not significant (**Supplementary Tables 2-4**).

**Figure 1.**
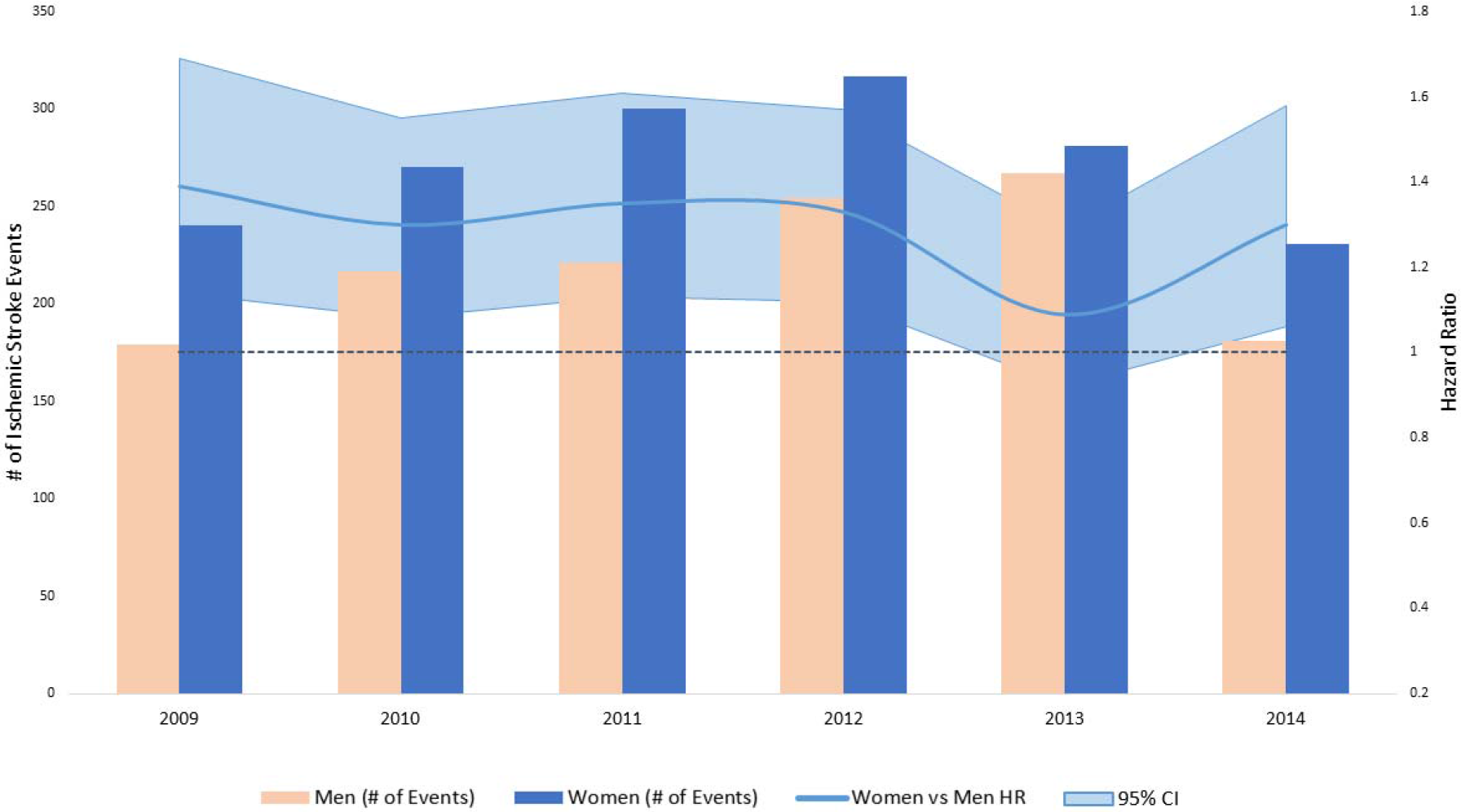
Population counts by sex and adjusted HRs (95% CI) for associations of sex with incidence of ischemic stroke in patients with atrial fibrillation by year, Optum Clinformatics® 2009-2015. HRs and 95% confidence intervals were calculated among atrial fibrillation (AF) patients from the Optum Clinformatics database using Cox proportional hazard modeling for the association between sex and ischemic stroke. The model was adjusted for age at AF diagnosis, race/ethnicity (black/white/Hispanic), education, and CHA2DS2-VASc scores, which assesses the risk of stroke among AF patients based on presence of congestive heart failure, hypertension, age above 75 years, previous stroke, vascular disease, diabetes, and age from 65 to 74. Stratification by year was based on year of AF diagnosis with a follow-up period of up to 365 days after date of AF diagnosis.

**Figure 2.**
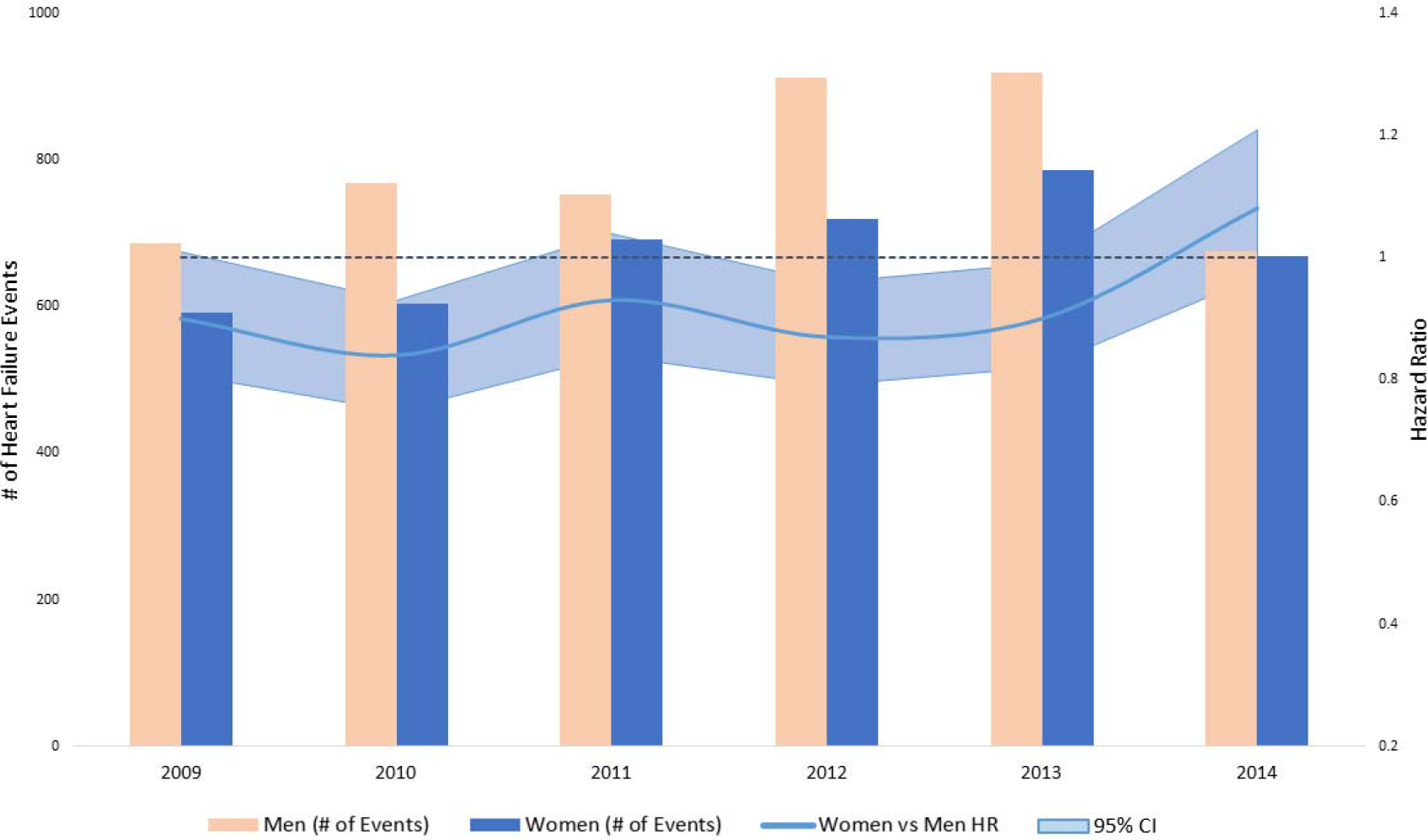
Population counts by sex and adjusted HRs (95% CI) for associations of sex with incidence of heart failure in patients with atrial fibrillation by year, Optum Clinformatics® 2009-2015. HRs and 95% confidence intervals were calculated among atrial fibrillation (AF) patients from the Optum Clinformatics database using Cox proportional hazard modeling for the association between sex and heart failure. The model was adjusted for age at AF diagnosis, race/ethnicity (black/white/Hispanic), education, and CHA2DS2-VASc scores, which assesses the risk of stroke among AF patients based on presence of congestive heart failure, hypertension, age above 75 years, previous stroke, vascular disease, diabetes, and age from 65 to 74. Stratification by year was based on year of AF diagnosis with a follow-up period of up to 365 days after date of AF diagnosis.

**Figure 3.**
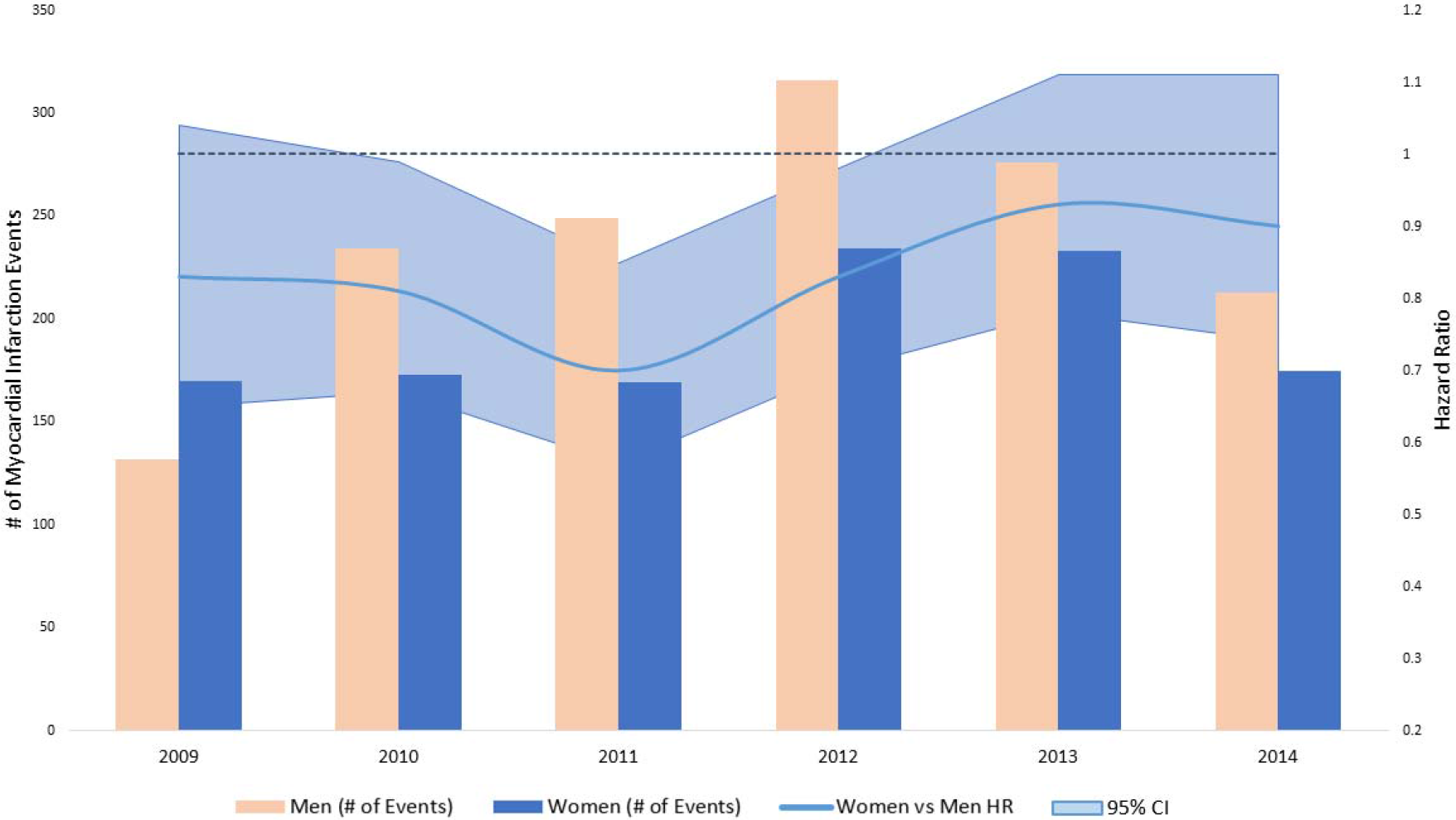
Population counts by sex and adjusted HRs (95% CI) for associations of sex with incidence of myocardial infarction in patients with atrial fibrillation by year, Optum Clinformatics® 2009-2015. HRs and 95% confidence intervals were calculated among atrial fibrillation (AF) patients from the Optum Clinformatics database using Cox proportional hazard modeling for the association between sex and myocardial infarction. The model was adjusted for age at AF diagnosis, race/ethnicity (black/white/Hispanic), education, and CHA2DS2-VASc scores, which assesses the risk of stroke among AF patients based on presence of congestive heart failure, hypertension, age above 75 years, previous stroke, vascular disease, diabetes, and age from 65 to 74. Stratification by year was based on year of AF diagnosis with a follow-up period of up to 365 days after date of AF diagnosis.

**Figure 4.**
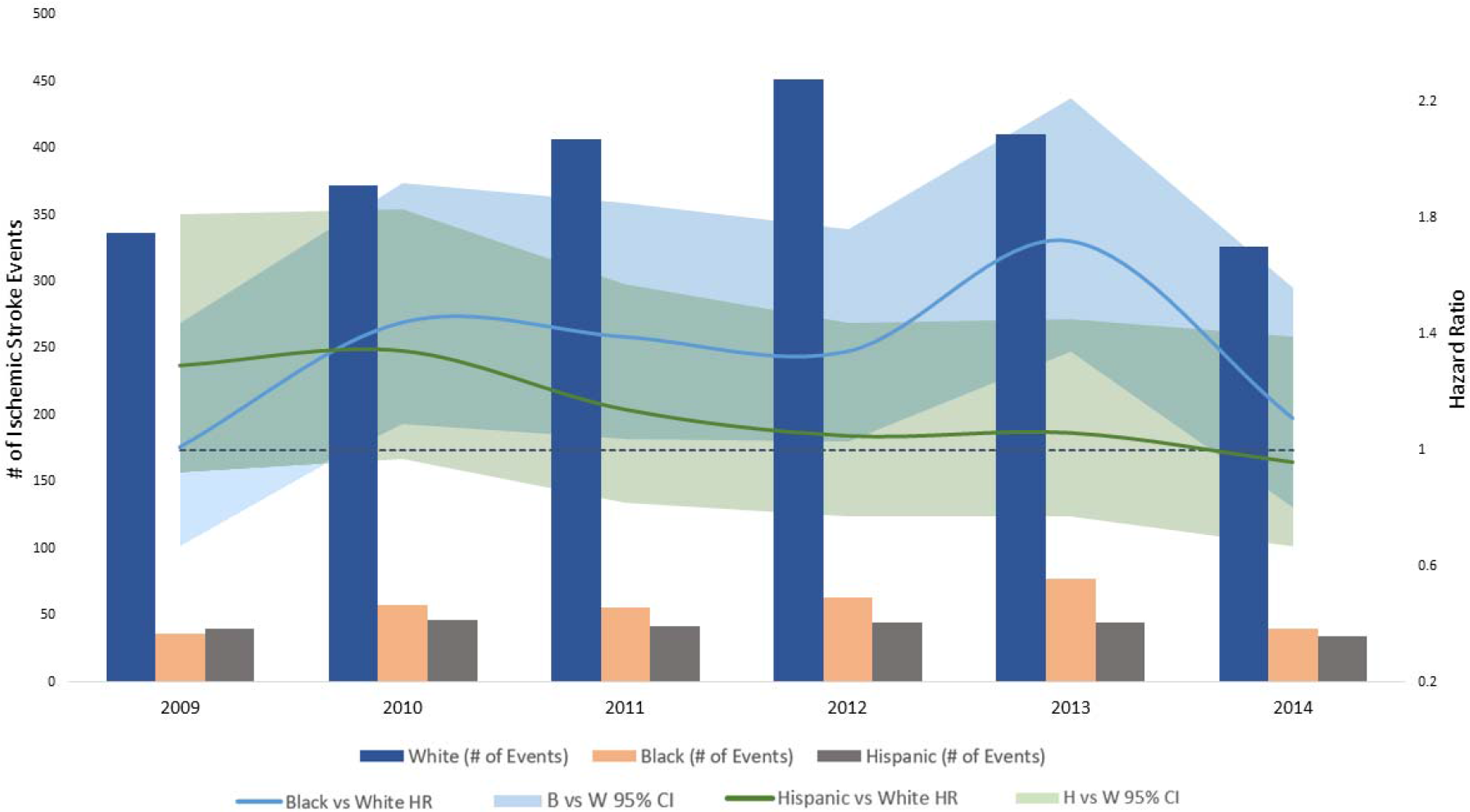
Population counts by race/ethnicity and adjusted HRs (95% CI) for associations of race/ethnicity with incidence of ischemic stroke in patients with atrial fibrillation by year, Optum Clinformatics® 2009-2015. HRs and 95% confidence intervals were calculated among atrial fibrillation (AF) patients from the Optum Clinformatics database using Cox proportional hazard modeling for the association between race/ethnicity and ischemic stroke. The model was adjusted for age at AF diagnosis, sex (female/male), education, and CHA2DS2-VASc scores, which assesses the risk of stroke among AF patients based on presence of congestive heart failure, hypertension, age above 75 years, previous stroke, vascular disease, diabetes, and age from 65 to 74. Stratification by year was based on year of AF diagnosis with a follow-up period of up to 365 days after date of AF diagnosis.

**Figure 5.**
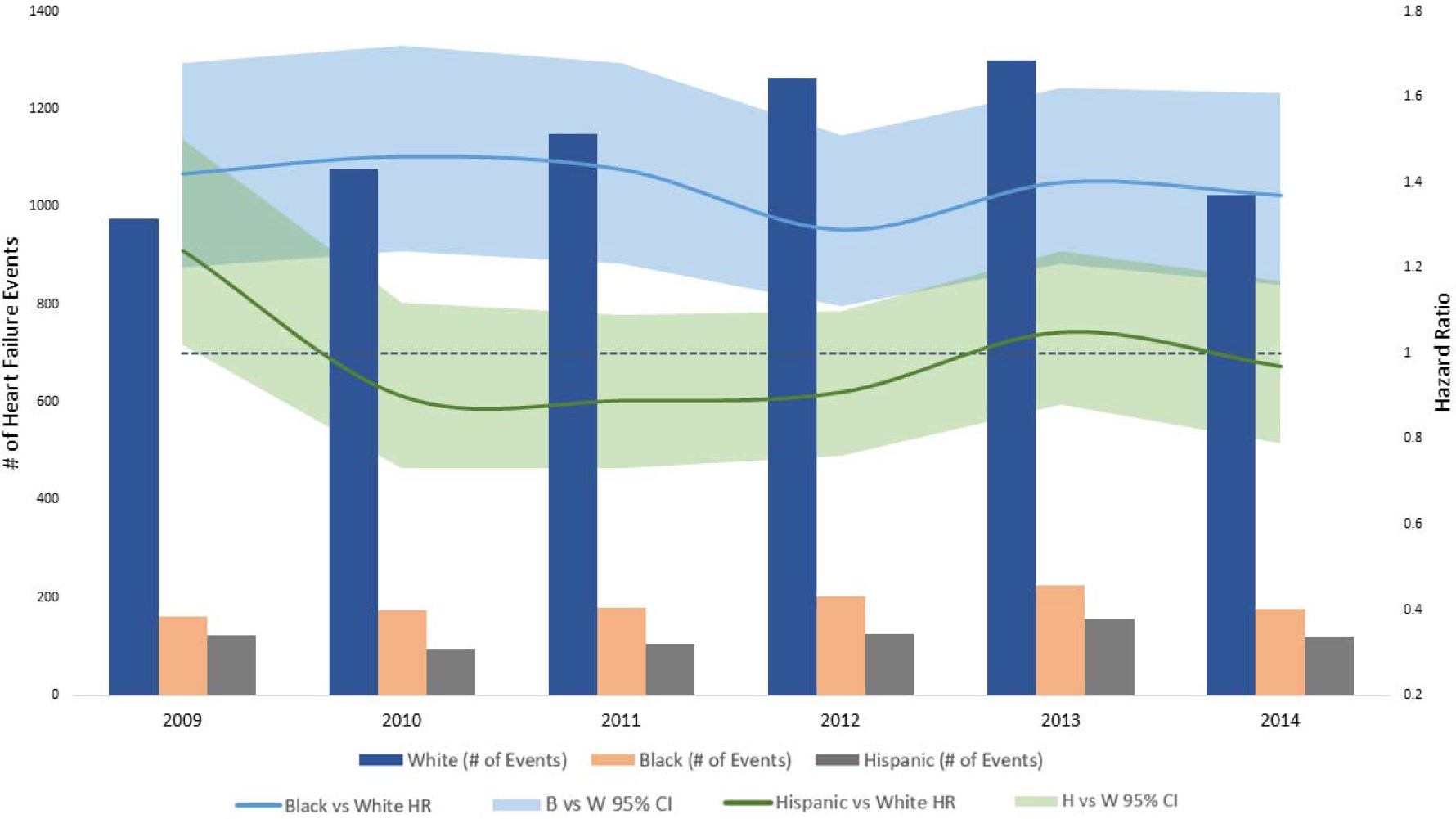
Population counts by race/ethnicity and adjusted HRs (95% CI) for associations of race/ethnicity with incidence of heart failure in patients with atrial fibrillation by year, Optum Clinformatics® 2009-2015. HRs and 95% confidence intervals were calculated among atrial fibrillation (AF) patients from the Optum Clinformatics database using Cox proportional hazard modeling for the association between race/ethnicity and heart failure. The model was adjusted for age at AF diagnosis, sex (female/male), education, and CHA2DS2-VASc scores, which assesses the risk of stroke among AF patients based on presence of congestive heart failure, hypertension, age above 75 years, previous stroke, vascular disease, diabetes, and age from 65 to 74. Stratification by year was based on year of AF diagnosis with a follow-up period of up to 365 days after date of AF diagnosis.

**Figure 6.**
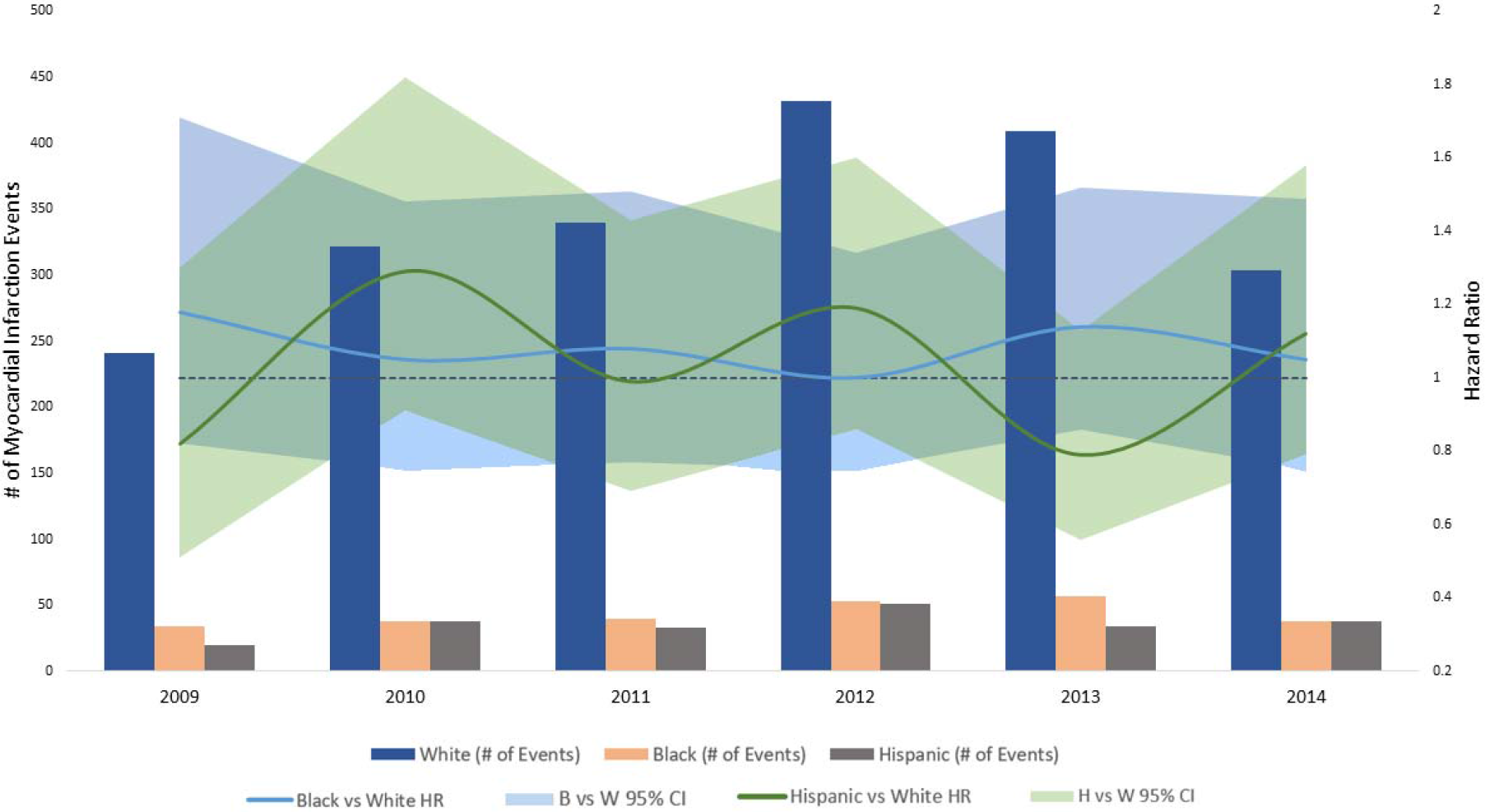
Population counts by race/ethnicity and adjusted HRs (95% CI) for associations of race/ethnicity with incidence of myocardial infarction in patients with atrial fibrillation by year, Optum Clinformatics® 2009-2015. HRs and 95% confidence intervals were calculated among atrial fibrillation (AF) patients from the Optum Clinformatics database using Cox proportional hazard modeling for the association between race/ethnicity and myocardial infarction. The model was adjusted for age at AF diagnosis, sex (female/male), education, and CHA2DS2-VASc scores, which assesses the risk of stroke among AF patients based on presence of congestive heart failure, hypertension, age above 75 years, previous stroke, vascular disease, diabetes, and age from 65 to 74. Stratification by year was based on year of AF diagnosis with a follow-up period of up to 365 days after date of AF diagnosis.

## DISCUSSION

In this large administrative claims database in the United States, women with AF experienced a higher rate of ischemic stroke compared with men, while men had a higher rate of heart failure and myocardial infarction. Black patients with AF had a higher rate of ischemic stroke, heart failure, and myocardial infarction compared to whites, while no differences were observed between whites, Hispanics, and Asian Americans. These patterns held steady in year-stratified analysis from 2009 to 2014, as shown by no statistically significant interactions between year of diagnosis and race/sex, indicating lack of evidence that these disparities have changed during this six-year period.

Recent reports have shown that women who have AF have a higher risk of ischemic stroke and myocardial infarction compared with men.^1^ Our results were consistent with an increased risk for stroke, but we did not observe a higher risk of myocardial infarction in women. In the general United States population, men are more likely to develop coronary heart disease compared with women,^7^ and this is also possibly true in our cohort of AF patients. Additionally, this may explain the higher risk for heart failure in men compared with women.

The data in this report also confirm that black patients with AF are more likely to develop cardiovascular disease events compared with whites. In the ARIC cohort, black patients with AF had a higher rate of stroke, heart failure, and coronary heart disease than whites.^2^ Our findings further confirm this black-white difference in a large claims database and demonstrate that the rates of these outcomes do not vary between whites and other races/ethnicities.

The last decade has experienced important changes in the ability to manage AF, with the approval of non-vitamin K antagonist oral anticoagulants (NOACs) for stroke prevention and the more common use of catheter ablation for AF rhythm control.^8, 9^ Unfortunately, management of AF—such as oral anticoagulation—is of lower quality in blacks compared to whites and in women compared to men in the US.^10, 11^ The lack of improvement in the disparities in risk of stroke and other cardiovascular outcomes suggests that previously described treatment disparities are not being properly addressed.

Racial/ethnic and sex disparities in NOAC use are evident, with women with AF being less likely to use NOACs than men, and black and Hispanic patients with AF using NOACs at nearly half the rate of their white counterparts.^12^ Total expenditure in the United States on NOACs increased from $95 million to $1.76 billion between 2010 and 2014 alone, with the out-of-pocket responsibility increasing from $119 million to $275 million during the same time period.^12^

These disparities also extend beyond medication. Between 2006 to 2011, women were less likely than men to undergo catheter ablation after hospital presentation with symptomatic AF, even though women experienced higher rates of AF-related rehospitalizations.^13^ Similarly, blacks and Hispanics were far less likely to undergo catheter ablation compared to whites, yet experienced higher rates of rehospitalization.^13^ Such gaps in management often result in insufficient prophylaxis for endpoints such as stroke and heart failure.

Our analysis should be interpreted in the context of its limitations. First, race is misclassified in these data. Second, both AF and cardiovascular outcomes were based on diagnostic codes included in healthcare claims, which may lead to misclassification. However, whenever possible, we used validated algorithms with high positive predictive value to define AF and endpoints. If we assume that the misclassification was non-differential, the true effect would, in expectation, be larger than the observed effect. Third, results may not be generalizable to populations without health insurance. Finally, other factors associated with race and sex may confound the reported associations with cardiovascular outcomes.

Overall, the findings in this report indicate the presence of sex heterogeneity in the rate of adverse cardiovascular outcomes in patients with AF, confirm the adverse risk profile in blacks compared with whites who have AF, and highlight lack of progress in reducing those differences. Further research is needed to understand these findings in order to develop targeted preventive strategies to improve outcomes in these subgroups of AF patients and reduce overall cardiovascular health disparities.

## Supporting information

Supplementary Results

## ACKNOWLEDGEMENTS

None.

