## Supplementary Results for "Sex and Racial Differences in Cardiovascular Disease Risk in Patients with Atrial Fibrillation"

**Supplementary Table 1. ICD-9-CM Codes for Covariates and Endpoint Diagnosis**

| **Outcome** | **ICD-9-CM Codes** |
| --- | --- |
| Diabetes mellitus | 250 |
| Ischemic Stroke | 433, 434, 435, 436, 437, 438 |
| Heart failure | 398.91, 402.01, 402.11, 402.91, 404.01, 404.03, 404.11, 404.13, 404.91, 404.93, 425.4, 425.9, 428 |
| Hypertension | 401, 402, 403, 404, 405 |
| Myocardial infarction | 410, 412 |
| Peripheral Artery Disease | 440.0, 440.2, 440.9, 443.9 |

**Supplementary Table 2. Year-stratified incidences of ischemic stroke across race/ethnicity and sex in patients with atrial fibrillation, Optum Clinformatics® 2009-2015**

|  | Men | Women | White | Black | Hispanic |
| --- | --- | --- | --- | --- | --- |
| **2009** |  |  |  |  |  |
| **N. Events** | 179 | 240 | 336 | 36 | 40 |
| **HR (95% CI)** | 1 (Ref) | 1.39 (1.14, 1.69) | 1 (Ref) | 1.01 (0.71, 1.44) | 1.29 (0.92, 1.81) |
| **2010** |  |  |  |  |  |
| **N. Events** | 217 | 270 | 372 | 58 | 47 |
| **HR (95% CI)** | 1 (Ref) | 1.30 (1.08, 1.55) | 1 (Ref) | 1.44 (1.09, 1.92) | 1.34 (0.97, 1.83) |
| **2011** |  |  |  |  |  |
| **N. Events** | 221 | 300 | 407 | 56 | 42 |
| **HR (95% CI)** | 1 (Ref) | 1.35 (1.13, 1.61) | 1 (Ref) | 1.39 (1.04, 1.85) | 1.14 (0.82, 1.57) |
| **2012** |  |  |  |  |  |
| **N. Events** | 254 | 317 | 451 | 63 | 45 |
| **HR (95% CI)** | 1 (Ref) | 1.33 (1.12, 1.57) | 1 (Ref) | 1.34 (1.03, 1.76) | 1.05 (0.77, 1.44) |
| **2013** |  |  |  |  |  |
| **N. Events** | 267 | 281 | 410 | 77 | 45 |
| **HR (95% CI)** | 1 (Ref) | 1.09 (0.92, 1.29) | 1 (Ref) | 1.72 (1.34, 2.21) | 1.06 (0.77, 1.45) |
| **2014** |  |  |  |  |  |
| **N. Events** | 181 | 231 | 326 | 40 | 34 |
| **HR (95% CI)** | 1 (Ref) | 1.30 (1.06, 1.58) | 1 (Ref) | 1.11 (0.80, 1.56) | 0.96 (0.67, 1.39) |
|  | Year-Sex Interaction | P = 0.17 | Year-Race Interaction | P = 0.31 |  |

HR, hazard ratio; CI, confidence interval.

^*^Cox model adjusted for age, sex, race/ethnicity, education and CHA_2_DS_2_-VASc.

**Supplementary Table 3. Year-stratified incidences of heart failure across race/ethnicity and sex in patients with atrial fibrillation, Optum Clinformatics® 2009-2015**

|  | Men | Women | White | Black | Hispanic |
| --- | --- | --- | --- | --- | --- |
| **2009** |  |  |  |  |  |
| **N. Events** | 686 | 592 | 975 | 162 | 122 |
| **HR (95% CI)** | 1 (Ref) | 0.90 (0.81, 1.01) | 1 (Ref) | 1.42 (1.20, 1.68) | 1.24 (1.02, 1.50) |
| **2010** |  |  |  |  |  |
| **N. Events** | 769 | 603 | 1076 | 174 | 94 |
| **HR (95% CI)** | 1 (Ref) | 0.84 (0.75, 0.93) | 1 (Ref) | 1.46 (1.24, 1.72) | 0.90 (0.73, 1.12) |
| **2011** |  |  |  |  |  |
| **N. Events** | 753 | 692 | 1149 | 179 | 104 |
| **HR (95% CI)** | 1 (Ref) | 0.93 (0.84, 1.04) | 1 (Ref) | 1.43 (1.21, 1.68) | 0.89 (0.73, 1.09) |
| **2012** |  |  |  |  |  |
| **N. Events** | 912 | 720 | 1264 | 201 | 125 |
| **HR (95% CI)** | 1 (Ref) | 0.87 (0.79, 0.96) | 1 (Ref) | 1.29 (1.11, 1.51) | 0.91 (0.76, 1.10) |
| **2013** |  |  |  |  |  |
| **N. Events** | 919 | 786 | 1298 | 226 | 155 |
| **HR (95% CI)** | 1 (Ref) | 0.90 (0.82, 0.99) | 1 (Ref) | 1.40 (1.21, 1.62) | 1.05 (0.88, 1.24) |
| **2014** |  |  |  |  |  |
| **N. Events** | 675 | 669 | 1023 | 176 | 119 |
| **HR (95% CI)** | 1 (Ref) | 1.08 (0.97, 1.21) | 1 (Ref) | 1.37 (1.16, 1.61) | 0.97 (0.79, 1.17) |
|  | Year-Sex Interaction | P = 0.07 | Year-Race Interaction | P = 0.89 |  |

HR, hazard ratio; CI, confidence interval.

^*^Cox model adjusted for age, sex, race/ethnicity, education and CHA_2_DS_2_-VASc.

**Supplementary Table 4. Year-stratified incidences of myocardial infarction across race/ethnicity and sex in patients with atrial fibrillation, Optum Clinformatics® 2009-2015**

|  | Men | Women | White | Black | Hispanic |
| --- | --- | --- | --- | --- | --- |
| **2009** |  |  |  |  |  |
| **N. Events** | 132 | 170 | 241 | 34 | 20 |
| **HR (95% CI)** | 1 (Ref) | 0.83 (0.65, 1.04) | 1 (Ref) | 1.18 (0.82, 1.71) | 0.82 (0.51, 1.30) |
| **2010** |  |  |  |  |  |
| **N. Events** | 234 | 173 | 322 | 38 | 38 |
| **HR (95% CI)** | 1 (Ref) | 0.81 (0.67, 0.99) | 1 (Ref) | 1.05 (0.75, 1.48) | 1.29 (0.91, 1.82) |
| **2011** |  |  |  |  |  |
| **N. Events** | 249 | 169 | 340 | 40 | 33 |
| **HR (95% CI)** | 1 (Ref) | 0.70 (0.57, 0.85) | 1 (Ref) | 1.08 (0.77, 1.51) | 0.99 (0.69, 1.43) |
| **2012** |  |  |  |  |  |
| **N. Events** | 316 | 234 | 432 | 53 | 51 |
| **HR (95% CI)** | 1 (Ref) | 0.83 (0.70, 0.98) | 1 (Ref) | 1.00 (0.75, 1.34) | 1.19 (0.86, 1.60) |
| **2013** |  |  |  |  |  |
| **N. Events** | 276 | 233 | 409 | 57 | 34 |
| **HR (95% CI)** | 1 (Ref) | 0.93 (0.78, 1.11) | 1 (Ref) | 1.14 (0.86, 1.52) | 0.79 (0.56, 1.13) |
| **2014** |  |  |  |  |  |
| **N. Events** | 213 | 175 | 304 | 38 | 38 |
| **HR (95% CI)** | 1 (Ref) | 0.90 (0.74, 1.11) | 1 (Ref) | 1.05 (0.75, 1.49) | 1.12 (0.79, 1.58) |
|  | Year-Sex Interaction | P = 0.34 | Year-Race Interaction | P = 0.82 |  |

HR, hazard ratio; CI, confidence interval.

^*^Cox model adjusted for age, sex, race/ethnicity, education and CHA_2_DS_2_-VASc.
